# Distinct routes to thermotolerance in the fungal pathogen *Cryptococcus neoformans*

**DOI:** 10.1101/2024.04.08.588590

**Authors:** Mara J. W. Schwiesow, Leah A. Farinella, Marina Ruzic, Jake T. Leinas, Nels C. Elde, Zoë A. Hilbert

## Abstract

Increasing temperatures associated with climate change have the potential for far-reaching impact on human health and disease vectors, including fungal pathogens. Pathogenic fungi occupy a wide range of environments across the world, and their ranges have been slowly expanding in recent decades due, in part, to climate change. Despite these links between increasing temperature and higher prevalence of fungal disease, the direct effects of rising environmental temperatures on the evolution of pathogenic fungi remains unclear. In this study, we investigated how increasing temperatures drive adaptive evolution in the human fungal pathogen *Cryptococcus neoformans*. First, we performed serial passages of a *C. neoformans* environmental isolate with gradual changes in temperature over the course of 38 days. Through this approach we identified several distinct thermally adapted isolates with competitive growth advantages over the parental strain at high temperatures. We then characterized the phenotypic and genetic changes acquired in these evolved isolates, which include alteration of cell size, colony morphology, and, notably, antifungal resistance. Our genetic analyses further revealed distinct genes that facilitate thermoadaptation in different populations—identifying new molecular players in the regulation of this trait and revealing that there are multiple independent routes to gaining thermotolerance. These results highlight the remarkable flexibility of fungi to adapt rapidly to new environments and raise pressing questions about the impacts of rising environmental temperatures on the future of infectious diseases and human health.

## Introduction

The impacts of planetary warming on biological systems hold pivotal implications for human and animal health worldwide^1^. Climate change is likely to have particularly dramatic effects in the realm of infectious disease, where increased environmental temperatures could fundamentally alter the number, nature, and geographic distribution of pathogenic microbes, including fungi^2^. However, little work has been done to investigate how incrementally increasing temperatures will affect the evolution of these pathogens and what consequences this adaptation may have in the context of human infection.

For fungi, in particular, the inability of most species to grow at the high body temperatures characteristic of mammals has long been considered a primary limiting factor in the ability of these microbes to establish infection^3^. As global temperatures rise, however, adaptation to higher temperatures is expected to drive increases in the number of fungal species capable of surviving at mammalian body temperature, perhaps even driving the emergence of new species with pathogenic potential^4^. Increased environmental temperature will also result in range expansions for pathogenic fungi. Indeed, this has already been observed in several fungal species including *Coccidioides* spp., *Cryptococcus gattii*, and *Candida auris*, with *C. auris* emerging on three separate continents simultaneously^5–7^. Although changes in the geographic range for these particular species are in part due to expansion of their preferred environments, there is also evidence to suggest that these organisms are adapting to new, more stressful environmental conditions^8,9^. Given these trends, it can be expected that as global temperatures continue to rise, the ranges of these and other pathogenic fungi will continue to expand, the threat of thermal adaptation in these organisms will escalate, and human body temperature may no longer pose such an impediment to fungal infection.

In this study, we focus on the role of increasing environmental temperatures in shaping the evolution of the common human fungal pathogen, *Cryptococcus neoformans*. *C. neoformans* is a pathogenic basidiomycete fungus found ubiquitously in the environment, usually residing in bird guano, soil, and trees, and infection by this species is currently estimated to cause over 100,000 deaths annually^10,11^. Strains of *C. neoformans* occupy a wide range of temperature environments having been isolated from all continents, excluding Antarctica, both from environmental and clinical settings^12–16^. Despite this, our understanding of how thermotolerance evolved in this species and may vary across strains is relatively limited.

Some studies of laboratory reference isolates have revealed candidate genes and pathways in *C. neoformans* that are important for success at high temperature. For example, heat shock proteins, and certain other conserved genetic pathways, such as *CDC11*, calcineurin, and the HOG pathway, all play roles in adaptation to the host environment, including adaptation to host temperature^17–21^. However, these studies largely focus on gene knockouts of the laboratory reference strain and sidestep a more realistic view of how this organism might adapt by modulating expression or function of these or other genes to survive under changing conditions.

Experimental evolution is a powerful tool to investigate the processes and mechanisms of adaptation to new environments in wide-ranging species. Studies of thermal adaptation, in particular, include the pioneering work of Bennett and Lenski, who used long-term experimental evolution to study the adaptive changes that occur in the bacterial species *Escherichia coli* under high temperature stress^22^. In fungal species, similar serial passaging approaches at consistent, high temperatures have proven powerful in generating heat-tolerant strains for use in industrial settings^23^. Additionally, two studies in the yeasts *Saccharomyces cerevisiae* and *Saccharomyces uvarum* have shown that gradual and stepwise temperature ramps can increase the thermotolerance of fungi by 1.5°C or more, suggesting that these fungi have a remarkable plasticity for tolerating temperature fluctuations^24,25^. However, studies exploring the thermal adaptation capabilities of environmental pathogenic fungi and related impacts on pathogenicity have been largely missing.

In this study, we leverage experimental evolution approaches to study the process of thermal adaptation in the clinically relevant *C. neoformans*. Through gradually increasing temperatures over many generations, we found that not only can *C. neoformans* adapt to produce genetically distinct strains with increased thermotolerance, but that this adaptation occurs on relatively short timescales. In only 38 days of continuous culturing, we observed increased thermotolerance of one environmental strain of *C. neoformans* characterized by improved growth at high temperature with clear fitness advantages over the parental strain. Thermal adaptation occurred across a range of media types and co-evolved with other phenotypic changes in cell size, colony morphology, and antifungal resistance. We further identify two genes, not previously known to regulate temperature sensitivity, which acquired mutations in distinct evolved populations and contribute to improved growth at high temperature. Our findings indicate that *C. neoformans* is capable of adapting to increasing global temperatures and provide a foundation on which to build an understanding of how our changing climate could impact human health and fungal disease.

## Results

### Variable thermotolerance across environmental isolates of *C. neoformans*

Environmental strains of *C. neoformans* inhabit diverse locations and are exposed to a variety of temperature ranges with varying nutrient availability. Although environmental isolates may generally experience more temperate climates, they are also likely subjected to wider temperature swings in contrast to clinical isolates, which are better able to grow at the high temperatures (37°C and above) of the human body. To test the thermotolerance properties of a panel of environmental *C. neoformans* isolates, we performed spot dilution assays and assessed the growth of these strains at two different temperatures: the standard laboratory culture temperature of 30°C and at a high temperature of 37°C, representative of human body temperature. To assess how thermotolerance might differ across different nutritional environments, we additionally assayed strains under rich (Yeast extract Peptone Dextrose [YPD]) and minimal (Yeast Nitrogen Base with glucose [YNB]) nutrient conditions. These assays revealed that thermotolerance varies widely, even between closely related strains (Figure 1). Some strains, such as Ftc217-1 and Jo278-1, grow well at high temperature, similar to the laboratory reference strain, H99 (Figure 1b and c). Others, like Muc387-1, exhibited moderate growth defects at high temperature. In general, growth differences at both temperatures were more pronounced on the minimal YNB media than on YPD.

**Figure 1.**
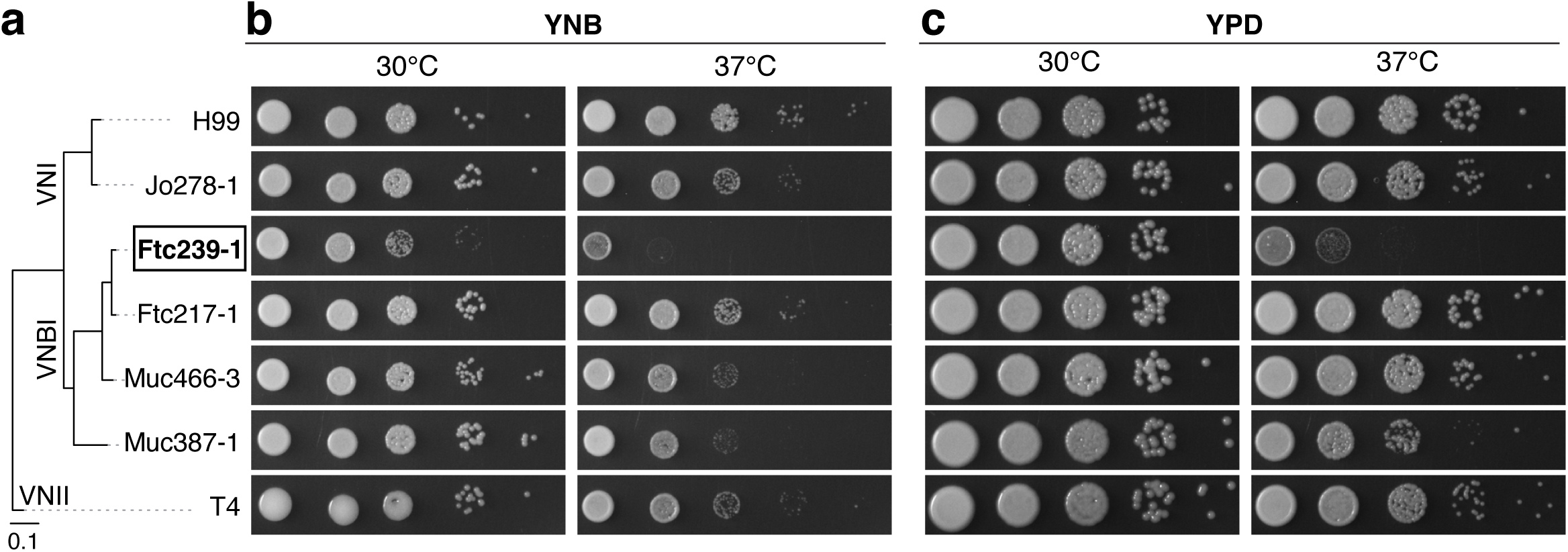
Phenotypic variation of *C. neoformans* environmental isolate growth at high temperatures. (a) Phylogenetic tree of *C. neoformans* strains used in this study adapted from Desjardins *et. al*. The primary strain used in subsequent figures, Ftc239-1, is indicated in bold and boxed. (b and c) Growth of *C. neoformans* strains on YNB (b) or YPD (c) agar plates at 30°C (left) or 37°C (right).

Among the set of strains tested here, one environmental isolate stood out: Ftc239-1. This strain, which was isolated from a mopane tree in Botswana, is closely related to Ftc217-1, but displays strikingly different growth across temperatures and nutrient conditions^14,15^. Unlike Ftc217-1, Ftc239-1 has severely attenuated growth at 37°C on both YNB and YPD. This strain also displays decreased growth at 30°C on YNB media (Figure 1b). The difference in growth on these two media types, across all strains tested but particularly in Ftc239-1, suggests that thermotolerance and nutrient conditions interface to control fungal growth.

Given the severe growth defects of Ftc239-1 at high temperatures, we reasoned that this would be an ideal strain to use to explore the mechanisms by which thermotolerance evolves in *C. neoformans*. In particular, the phenotypes of this strain gave us an opportunity to explore how it might adapt to higher temperatures relevant for human infection, while also providing an avenue to investigate the role of nutrient conditions in thermal adaptation.

### Experimental evolution can drive fungal adaptation to increasing temperatures

We used laboratory-based experimental evolution to study the process of thermal adaptation in the Ftc239-1 *C. neoformans* strain. Although temperature increases associated with human-induced climate change are rapid for geologic time, they are much more gradual for the organisms experiencing those changes. As such, we designed our experimental evolution paradigm to mimic the gradual temperature increases environmental pathogens may experience in the coming decades, though over a greatly abbreviated timescale. Specifically, Ftc239-1 was serially passaged every 48 hours starting at 30°C with temperature increases of 0.5–1°C every 4–6 days until a temperature of 35°C was reached (Figure 2a). In total, the fungal cells were grown continuously in this paradigm for 19 passages (38 days), which we estimate represents <150 generations of growth—a relatively short course of evolution. From the originating Ftc239-1 strain, eight populations were evolved in parallel, with four passaged in a rich media environment (YPD) and four passaged under nutrient-limiting conditions (YNB). Given the differences we observed in growth across these two media types between strains (Figure 1), along with previous work highlighting the role of several metabolic pathways in thermal sensitivity, we reasoned that the differing nutrient conditions would reveal distinct evolutionary paths^26,27^. These independently evolving <u>T</u>hermal <u>A</u>daptation populations were named TA1-4 for the populations evolved in YPD and TA5-8 for the YNB-passaged replicates (Figure 2a).

**Figure 2.**
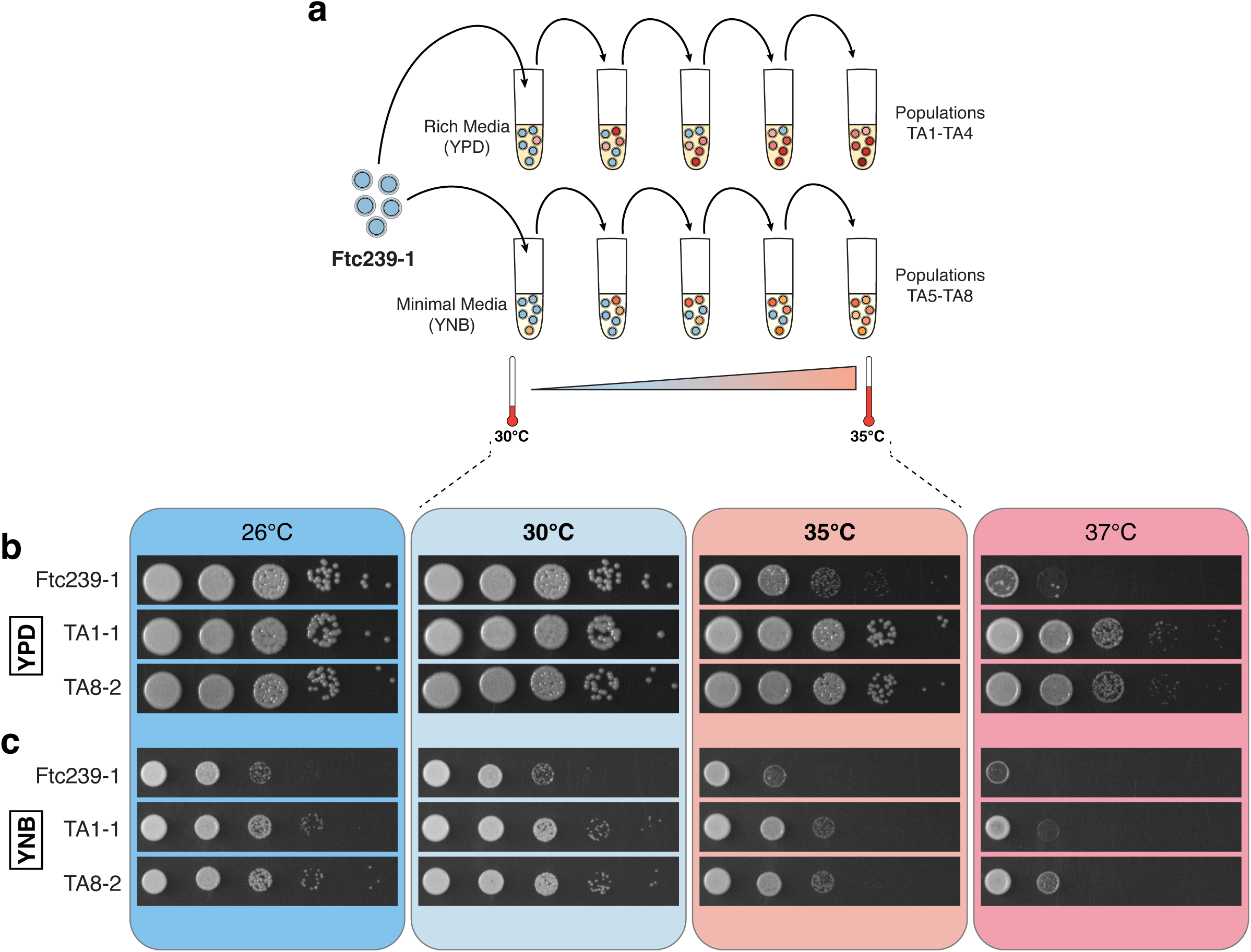
Serial passaging of *C. neoformans* selects for strains with increased thermotolerance. (a) Schematic of the serial passaging set-up used in this study. See also Materials and Methods. Populations TA1-TA4 were passaged in YPD; TA5-TA8 were passaged in YNB. (b and c) Growth of serially passaged isolates on YPD (b) or YNB (c) agar plates. Strains were tested for growth at temperatures they experienced during the experimental evolution passaging (30 and 35°C, middle panels). Growth was also assessed at temperatures outside of the range used during passaging (26°C and 37°C, outer panels).

All of the populations were viable until the end of the evolution experiment (35°C), indicating that this *C. neoformans* strain has the ability to adapt to or at least withstand increased thermal stress. Following the end of passaging, we harvested and froze the evolved populations; we also recovered individual clonal isolates for further characterization (see Methods). Intriguingly, we observed variation in the appearances and morphology of the clonal samples at the end of the evolution. Some displayed a mucoid appearance, while others had increased colony size compared to the starting isolate, providing our first indication that our experimental evolution had yielded new and potentially adaptive phenotypes in these populations (Tables S1 and S2). Further, changes in these phenotypes were not always universal within populations, suggesting that there was both phenotypic and genetic heterogeneity at the end of our course of evolution.

### Experimentally evolved isolates grow better on solid media at high temperatures

Following passaging, we first set out to determine whether any of the individual endpoint colonies displayed increased growth at high temperature compared to the parental strain. We, again, employed spot dilution assays to assess the growth phenotypes of our 36 recovered clonal isolates (Figure S1). This revealed that every independently evolved population contained fungi with enhanced growth at 35°C compared to the parental. These spot assays also confirmed our previous observations that there was phenotypic heterogeneity across isolates both between populations and within a single population. This was particularly clear among YPD-evolved isolates from populations TA1-TA4 where individual isolates varied both in ability to grow at high temperature (compare TA2-3 and 2-5 to other TA2 isolates) and in ability to grow on YNB minimal media (see isolates TA3-1, TA3-4, and all isolates from the TA4 population). These data indicate that there are multiple distinct paths to gaining thermotolerance in this *C. neoformans* strain.

From these experiments, we selected individual isolates from two independent populations for further characterization and follow-up (Figure 2b and c). One of these isolates was evolved in YPD (TA1-1) and one was evolved in YNB (TA8-2). At 30°C, both TA1-1 and TA8-2 had increased growth on YNB, while maintaining similar growth on YPD to the parental strain (Figure 2b and c, inner panels). At 35°C, both evolved isolates grew one or two dilutions further than the parental strain. This effect was consistent on both media types suggesting that thermotolerance in these particular isolates confers an advantage across nutrient conditions.

We next tested whether the increased thermotolerance in TA1-1 and TA8-2 conferred an enhanced ability to grow at high temperatures beyond the range of our evolution and whether there are tradeoffs for increased thermotolerance, perhaps affecting growth at temperatures below 30°C. Thermal adaptation studies with other microbial species, like *E. coli*, have shown that such paradigms can select for evolved isolates with highly specific temperature preferences i.e. optimized for growth at the temperatures using during the experiment^22^. To assess the growth phenotypes of our strains outside the bounds of the experimental evolution, we looked at growth of our selected isolates in parallel at 26, 30, 35 and 37°C. Both thermally-adapted isolates showed increased growth at 37°C on both YPD and YNB medias (Figure 2b and c, outer panels). There was no substantial difference in growth from the parental strain for either isolate at 26°C on YPD, however there was a slight increase in growth of the evolved strains on YNB at this temperature. These results suggest that the gradual adaptation of this *C. neoformans* strain to a specific high temperature (35°C) confers a growth advantage in high temperatures generally, with no apparent tradeoffs for growth at lower temperatures.

### Thermally adapted strains outcompete the parental strain at 35°C

To directly test whether the evolved strains gained a fitness advantage for growth at high temperature, we conducted co-culture competition experiments. In these competition experiments, evolved strains were grown together with a drug-resistant version of the parental strain in the same liquid media in which they were originally evolved (Figure 3a).

**Figure 3.**
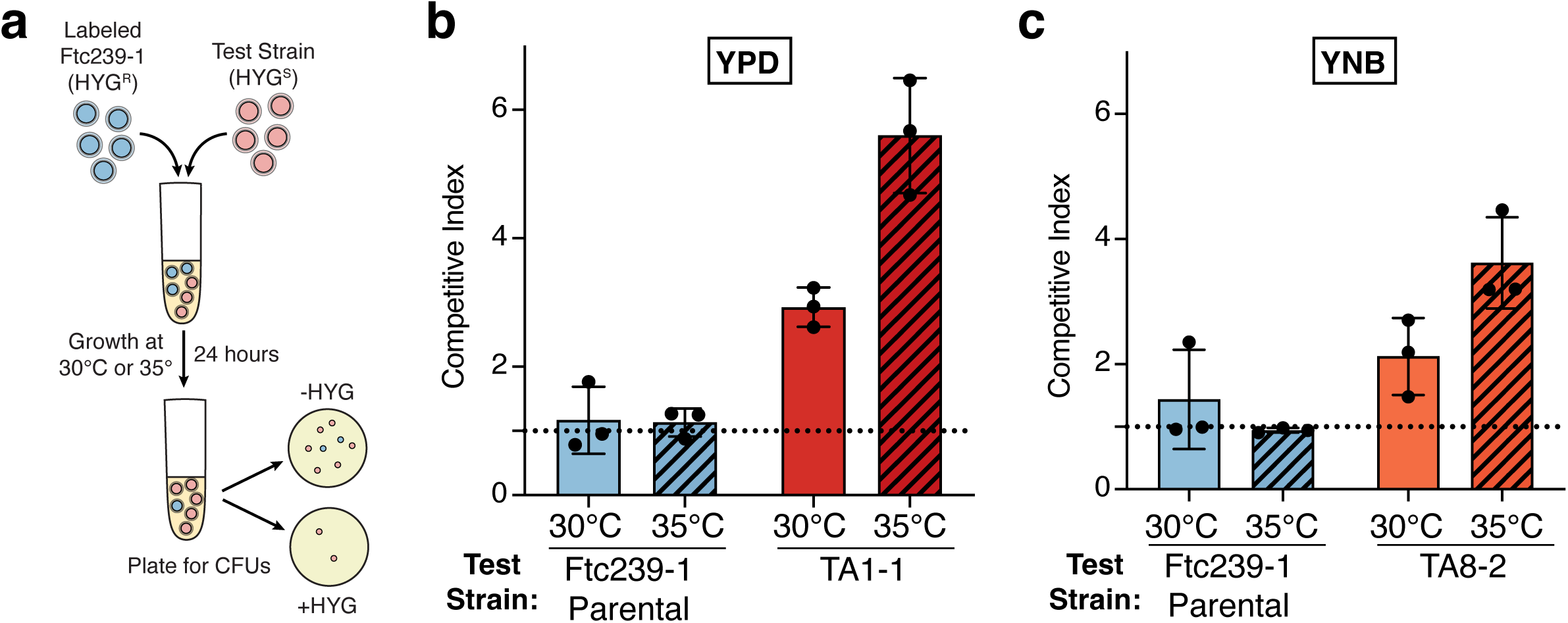
Thermally-adapted isolates outcompete the parental strain at high temperature. (a) Schematic of the competition experiment set-up between the HYG^R^-labeled parental strain and serially-passaged isolates at 30°C or 35°C. See also Materials and Methods. (b and c) Competition experiments between HYG^R^-labeled parental and unlabeled parental (blue), TA1-1 (dark red), TA8-2 (orange) strains in either YPD (b) or YNB (c) at both 30°C and 35°C. Competitive indices (CIs) are calculated as indicated in Methods. A CI=1 (dotted line) indicates equal growth of the two strains. Plotted values indicate the average value ± SEM of three replicate experiments carried out on three separate days; dots indicate the average value of biological replicates from a single experiment.

These experiments confirmed that each of the evolved strains has a substantial competitive advantage over the parental strain at 35°C. TA1-1 grew over five times better than the parental strain, and TA8-2 had a three to four-fold advantage in the media conditions used during the evolution experiment (Figure 3b and c). While this fitness effect is less robust at 30°C, the competitive advantages of the evolved strains were maintained at lower temperatures as well. These data indicate that the evolved isolates have successfully adapted to be able to grow at high temperatures (35°C) and further reveal, that these isolates have a fitness advantage over the parental strain not only at high temperature but across a range of temperatures.

### Evolved isolates have altered colony morphologies and titan cell production

We next asked whether thermal adaptation in our evolved strains was accompanied by alteration of other physical or morphological properties of the fungi. High temperature is a known regulator of different aspects of cryptococcal biology, including pathogenicity-related phenotypes such as capsule production, cell size, and melanization, among others^28^. Investigation of colony morphology revealed differences in our harvested clonal samples. The parental strain, Ftc239-1, has smooth, small colonies that are cream-white in color. The evolved strains (TA1-1 and TA8-2) exhibited changes in one or more of these features. TA1-1 colonies were consistently larger than the parental colonies and took on a mucoid appearance (Figure 2b, Table S1). These mucoid colonies have a glossy appearance with smooth edges and often form a looser pellet when spun down in liquid culture^29^. TA8-2 maintained the smooth colony phenotype seen in the parental strain, but the colonies were markedly larger (Table S2). Interestingly, more than half of the remaining endpoint colonies appeared mucoid, which suggests that the mucoid phenotype confers some advantage for surviving thermal stress or that there may be overlap in the genetic pathways that control these two traits (Tables S1 and S2). We hypothesized that changes in mucoidy might indicate a perturbation of the cell surface of these evolved cells, including production of the polysaccharide capsule that serves as a protective coating on cryptococcal cells under host-like conditions. However, our preliminary analysis of capsule production in our selected clonal isolates showed no major differences in capsule size (Figures S2a and b). This suggests that the mucoidy we observe in isolates across our evolved populations is likely derived from other altered features of our thermoadapted strains.

Given that colony morphology varied across our evolved isolates, we were curious what other aspects of cell morphology might be altered in these strains. To this end, we looked at the ability of our evolved isolates to form titan cells: large (>10 µm), polyploid cells that are induced during *in vivo* infection and *in vitro* under host-like conditions, including elevated temperatures^30^. Larger cell sizes inhibit phagocytosis by immune cells and a polyploid genome enables increased mutational flexibility, both of which allow the yeast to survive and adapt to host conditions during infection. Using an *in vitro* titanization assay, we observed substantial changes in titan cell formation in our selected thermoadapted isolates. Although TA1-1 showed robust titan cell formation, the TA8-2 strain failed to titanize entirely (Figures S2a and c). A second clonal isolate from the TA8 population, TA8-3, similarly failed to titanize under *in vitro* inducing conditions, suggesting that thermal adaptation in the TA8 population was accompanied by a cost for cell size variation (Figure S2C).

The opposing titanization phenotypes of TA1-1 and TA8-2 led us to ask whether titan cell formation was preserved in our other thermally adapted populations and if there might be a correlation between alterations in this phenotype and the media conditions each population experienced during the evolution. To assess this, we repeated our titan cell assays with 12 additional endpoint colonies with representatives from every independent population (Figure 4a). The majority of the isolates we examined in these experiments showed increased growth at high temperature compared to the parental starting strain (Figure S1). However, two isolates, TA2-5 (from YPD-evolved population TA2) and TA6-2 (from YNB-evolved population TA6) display parental-like temperature sensitive growth phenotypes and were included as important reference points for correlating cell morphology phenotypes with thermoadaptation.

**Figure 4.**
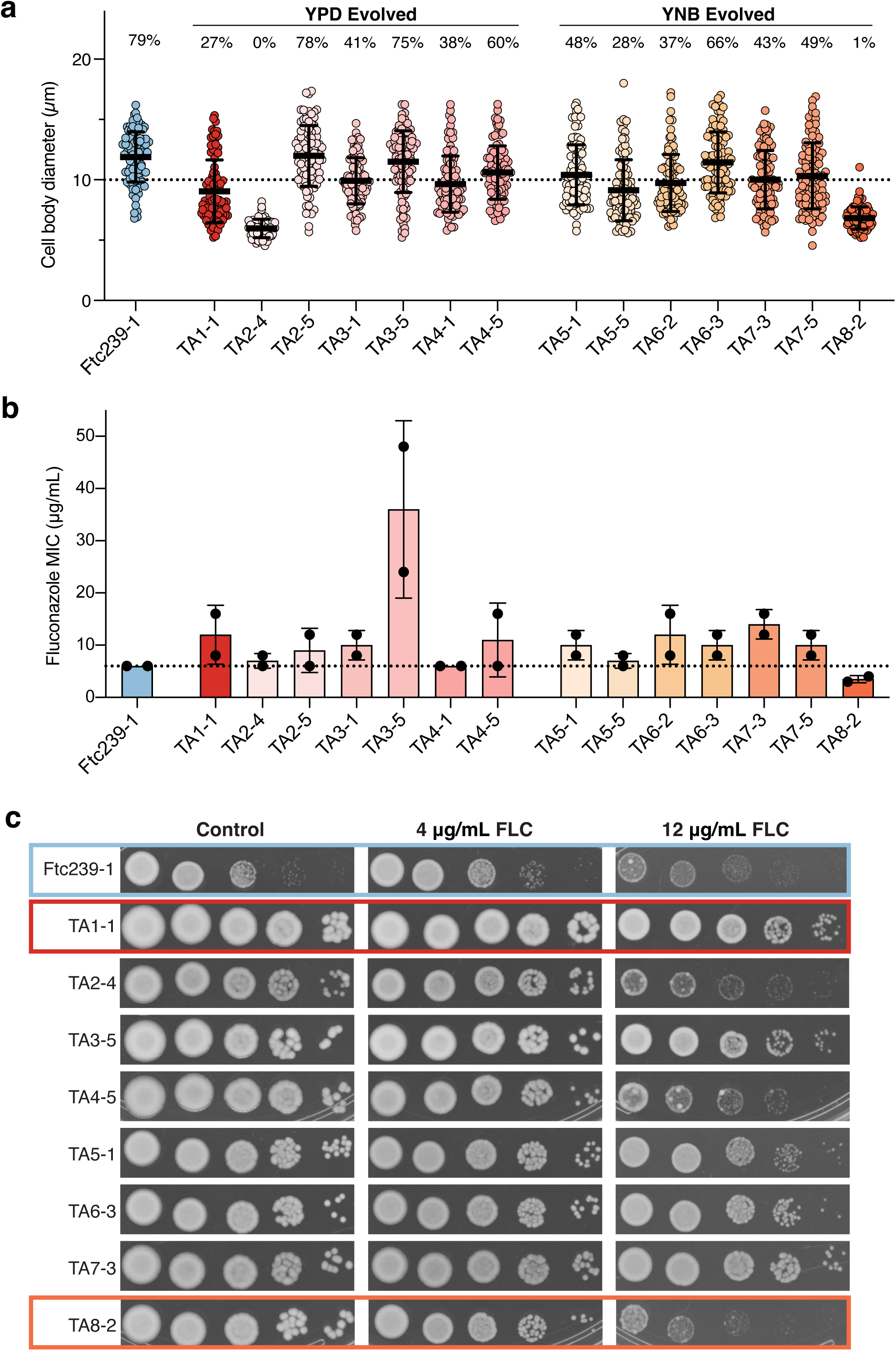
Thermal adaptation alters titan cell formation and antifungal drug susceptibility in serially-passaged isolates. (a) Measurements of cell body diameter from cells grown under titan inducing conditions. Percentages above each strain indicate the percent of cells with a diameter >10 µm (dotted line). Error bars show mean ± SD for one representative experiment. 100 cells per strain were measured with the exception of TA2-4 which displayed a growth defect under these conditions and only 60 cells could be quantified. (b) MIC values for expanded panel of clonal isolates as determined using fluconazole E-tests. Bars show the mean and ± SD of two replicate experiments with individual values plotted as dots. Mean MIC numerical values are also provided in Table 1. (c) Growth of a subset of isolates on control YPD plates, or YPD plates containing 4 or 12 µg/mL of the antifungal fluconazole. Images show 72 hours of growth at 35°C. The parental strain along with the two main isolates examined are boxed in color.

**Table 1.** Average fluconazole MIC values for experimentally evolved isolates. Numerical values of the data plotted in Figure 4b. Averages are for two MIC readings taken from different experimental replicates (on different days).

| Strain | Evolution Media | Growth at 37°C | Average FLC MIC (µg/mL) |
| --- | --- | --- | --- |
| Ftc239-1 | -- | - | 6 |
| TA1-1 | YPD | + | 12 |
| TA2-4 | YPD | + | 7 |
| TA2-5 | YPD | - | 9 |
| TA3-1 | YPD | + | 10 |
| TA3-5 | YPD | + | 36 |
| TA4-1 | YPD | + | 6 |
| TA4-5 | YPD | + | 11 |
| TA5-1 | YNB | + | 10 |
| TA5-5 | YNB | + | 7 |
| TA6-2 | YNB | - | 12 |
| TA6-3 | YNB | + | 10 |
| TA7-3 | YNB | + | 14 |
| TA7-5 | YNB | + | 10 |
| TA8-2 | YNB | + | 3.5 |

Although there was variation in the percentage of cells that exceeded the 10 µm threshold, we observed that the vast majority of the isolates from our evolved populations maintained the ability to titanize (Figure 4a). Besides TA8-2, only one other isolate failed to produce cells larger than 10 µm: the YPD-evolved TA2-4. Interestingly, the TA2-5 isolate from the same population, which shows impaired growth at high temperature, was still able to form titan cells, suggesting that the lack of titanization in TA2-4 (and other strains) may be a direct tradeoff caused by high temperature adaptation. Collectively, our investigations of cell morphology across our evolved isolates showt that these features are variably altered across and within independently evolved populations, regardless of media type, and include loss of key virulence-related traits like titanization.

### Antifungal drug susceptibility is altered in thermally adapted strains

In addition to thermal stress, fungal cells are subjected to a wide range of other stressors both in the environment and during infection, prompting us to ask whether our evolved isolates may also be more resistant to other types of cellular stress. We decided to explore this question using exposure to the antifungal drug fluconazole as a clinically relevant test to investigate the interface between thermal adaptation and other possible stressors.

In preliminary experiments assessing the minimum inhibitory concentrations (MICs) of our strains, we observed that our selected clonal isolates showed opposite trends in their MICs compared to the parental strain (Figure S2d). TA1-1 displayed enhanced fluconazole resistance compared to the parental strain and TA8-2 showing decreased resistance (Figure S2d). To gain a more comprehensive view of antifungal resistance in our thermally adapted strains, we assessed the MICs across our wider panel of isolates (Figure 4b). Strikingly, we observed that many strains displayed increased MICs compared to the parental Ftc239-1 strain, with a few notable strains (e.g. TA3-5) showing robust resistance to fluconazole (Figure 4b and Table 1). Only a single isolate from the evolution experiment, TA8-2, had a markedly decreased MIC in these experiments.

To gain a more nuanced view of fluconazole’s effects on growth in these isolates, we used spot dilution assays to observe growth on fixed concentrations of fluconazole (Figure 4c and Figure S3). We observed very few changes in growth for most strains on plates containing a low (4 µg/mL) dose of fluconazole; in contrast, high concentrations (12 µg/mL) of fluconazole triggered a much more variable growth inhibition. Although the parental strain and several others display marked decreases in growth when exposed to this concentration of fluconazole, many of the thermally adapted isolates continued to grow robustly showing that the increased MIC values are indeed indicative of increased antifungal resistance. This includes the isolates: TA1-1, TA3-5, TA5-1, TA6-3, and both isolates from the TA7 population.TA8-2, in contrast was consistently among the worst performers in these fluconazole assays, aligning with its lower MIC values. Interestingly, we observed that our isolates that remained temperature sensitive after passaging (TA2-5 and TA6-2) showed similar sensitivity to fluconazole as the parental strain, suggesting that drug resistance correlates with thermoadaptation in many of our experimentally evolved populations.

### Evolved isolates have distinct mutations that vary between populations and drive the emergence of thermotolerance

Given the strong growth advantage of our passaged strains along with the phenotypic differences across these isolates, we hypothesized that they likely carried adaptive genetic lesions that underlie their phenotypes. To begin to assess the genetic basis of adaptation in our thermally-adapted isolates, we returned our focus to TA1-1 and TA8-2, which differ in most traits beyond thermoadaptation, suggesting that they may have taken different evolutionary and genetic routes to adapt to high temperature. We performed whole-genome sequencing of our parental isolate, TA1-1, and TA8-2, along with additional isolates from the TA8 population.

We observed unique mutations in colonies from the same population (TA8-2 vs. other TA8 isolates) as well as across populations (YPD vs YNB, TA1-1 vs TA8-2). In the YPD-passaged TA1-1, we identified four fixed mutations affecting a range of different genes and with varying effects, from intronic mutations to frameshifting insertions (Table 2). In the TA8 YNB-evolved population, we identified six mutations that were shared across all clonal isolates, including TA8-2 (Table 3). These mutations were also at high frequency (>90%) in sequencing data from the pooled population culture indicating they were fixed in the TA8 population. In addition to the six mutations that were shared, we also identified several additional mutations that were present in some—but not all—clonal isolates in the TA8 population (Table 4). Notably, TA8-2 had no mutations beyond those that were fixed in the population. As with TA1-1 the identified mutations in TA8-2 and other TA8 isolates were dispersed across a number of genes and genomic locations and had a range of predicted effects on protein function.

**Table 2.**
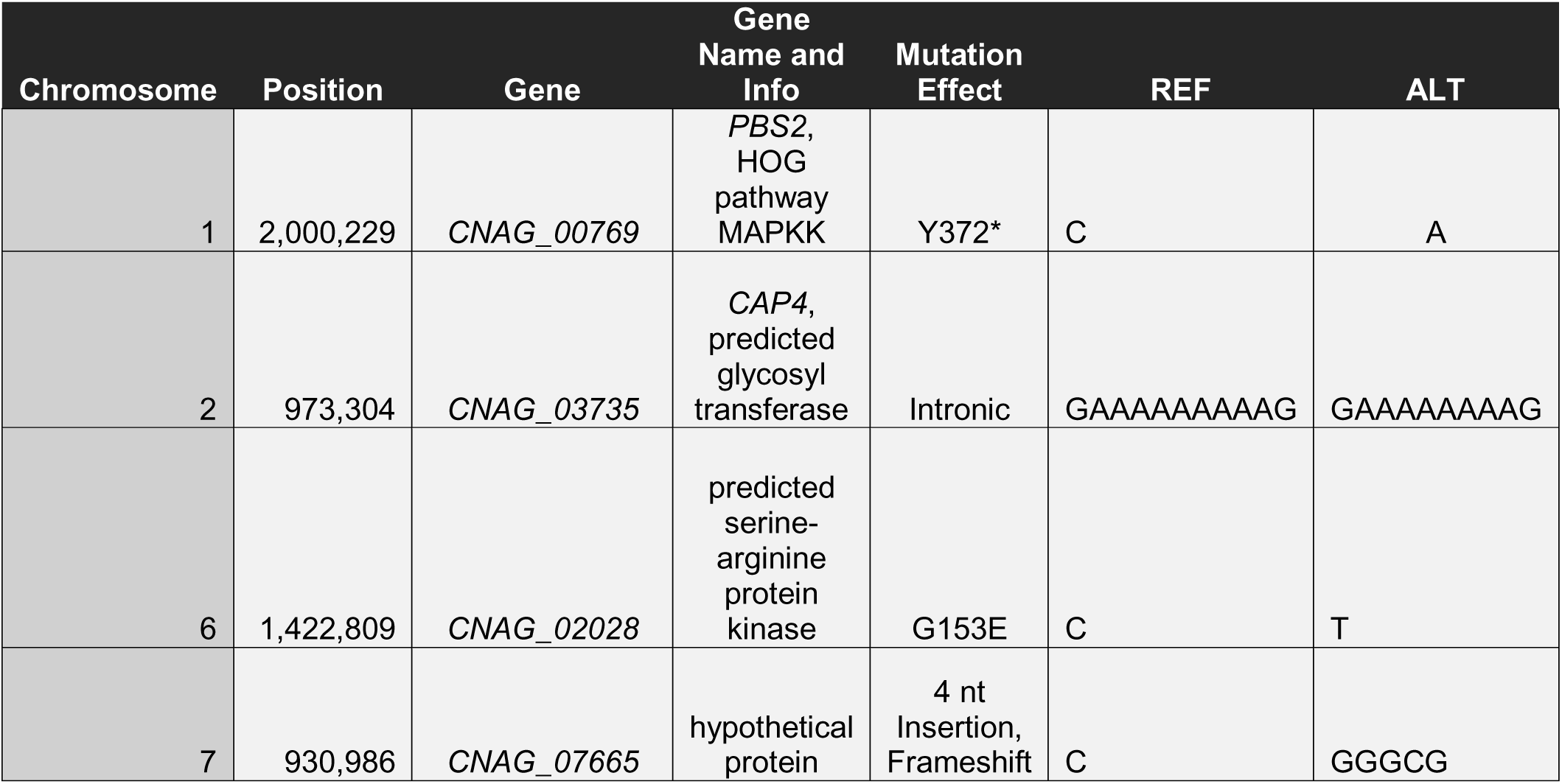
Fixed mutations in the TA1-1 YPD-passaged strain.

**Table 3.** Fixed mutations in the TA8 YNB-passaged population. These mutations are present in all of the sequenced TA8 clonal isolates.

| Chromosome | Position | Gene | Gene Name and Info | Mutation Effect | REF | ALT |
| --- | --- | --- | --- | --- | --- | --- |
| 2 | 1,592,272 | CNAG_06765 | LMP1, low mating performance protein | 1 nt deletion, Frameshift | TT | T |
| 4 | 328,289 | CNAG_05063 | SSK2, HOG pathway MAPKKK | 1 nt deletion, Frameshift | TT | T |
| 5 | 723,192 | Intergenic |  |  | G | C |
| 8 | 435,015 | CNAG_03438 | HXT1, hexose transporter | G63R | G | C |
| 10 | 1,058,846 | Intergenic (CNAG_04922 3' UTR) | CNAG_04922, hypothetical protein |  | C | A |
| 14 | 928,714 | Intergenic | Intergenic |  | C | T |

**Table 4.**
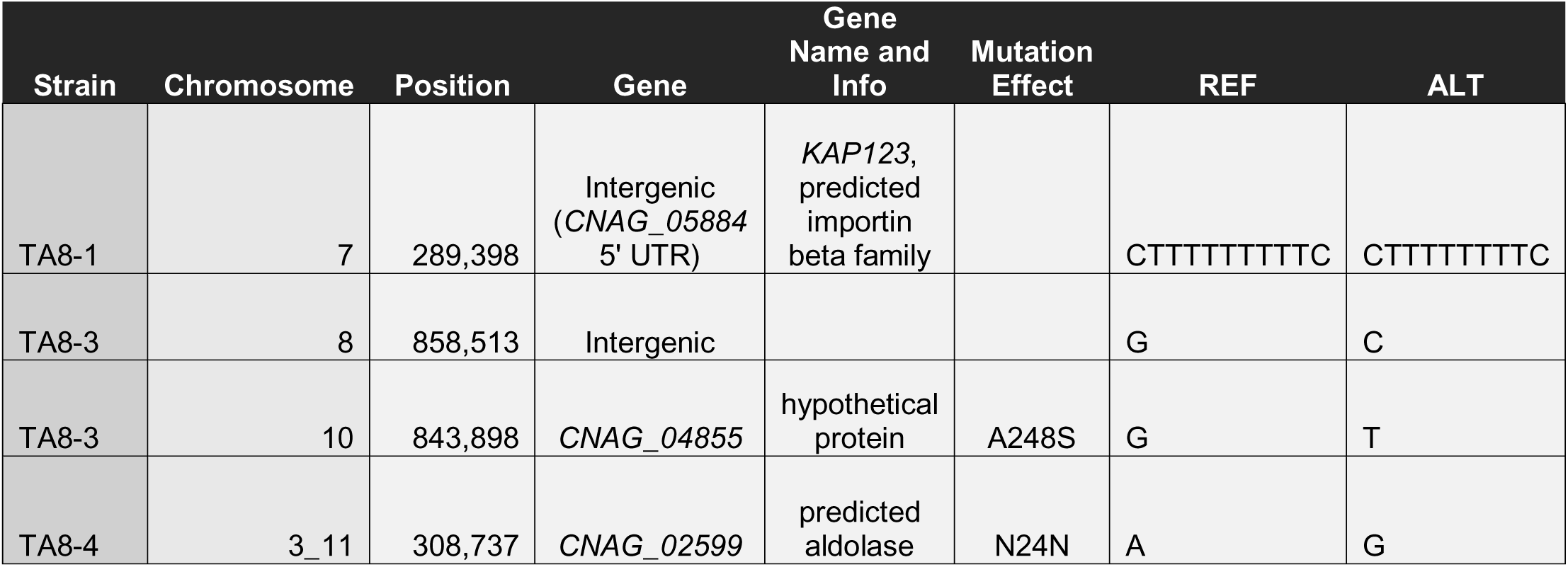
Additional fixed mutations in TA8-derived strains. These mutations are only present in the indicated colony from the TA8 population.

Although there was no overlap in the mutated genes between TA1-1 and TA8-2, we did note that both independent populations harbored mutations in core genes of the HOG pathway, which regulates cellular stress responses in *C. neoformans* and other related species^31^.In particular, we identified a missense mutation in the *PBS2* gene of the TA1-1 isolate and a frameshift mutation in the gene *SSK2* in the TA8 population strains^32^. The fact that we recovered mutations in two key HOG-pathway genes in these independently evolved populations highlights the importance of this pathway in responding to environmental stress and led us to hypothesize that the HOG pathway may play an important role in regulating temperature adaptation in this experiment.

To test this hypothesis and further probe the genetic basis of thermoadaptation in the TA1-1 and TA8-2 isolates, we constructed a series of gene deletion mutants, focusing on genes where we recovered coding sequence mutations. Spot assays of these mutants revealed that deletion of the HOG pathway genes, *PBS2* and *SSK2*, had no impact on growth at high temperature (Figure 5). Deletion of the genes *CNAG_07665* and *LMP1,* which carry frameshifting mutations in TA1-1 and TA8-2, respectively, similarly had no impact on growth at high temperature. Rather, we found that deletion of either the hexose transporter, *HXT1* (CNAG_03438), or an unnamed, uncharacterized predicted serine-arginine protein kinase (CNAG_02028) were sufficient for increased growth at both 35°C and 37°C when introduced to the Ftc239-1 parental strain. The high temperature growth phenotypes of these deletion mutants looked remarkably similar to those in our evolved clonal isolates, TA1-1 and TA8-2 which acquired mutations in these genes via experimental evolution (Figure 5 and Tables 2 and 3). These results strongly suggest that thermotolerance evolved through distinct mechanisms in these two replicate populations: with mutation of *CNAG_02028* driving this trait in the TA1 population and an *HXT1* lesion promoting thermotolerance in TA8. Although it remains to be seen what the impacts of the specific recovered mutations are on thermal adaptation along with the molecular mechanisms by which these genes contribute to thermotolerance, these results highlight the power of this experimental platform to uncover new understanding of the biology of this critically important trait in fungi.

**Figure 5.**
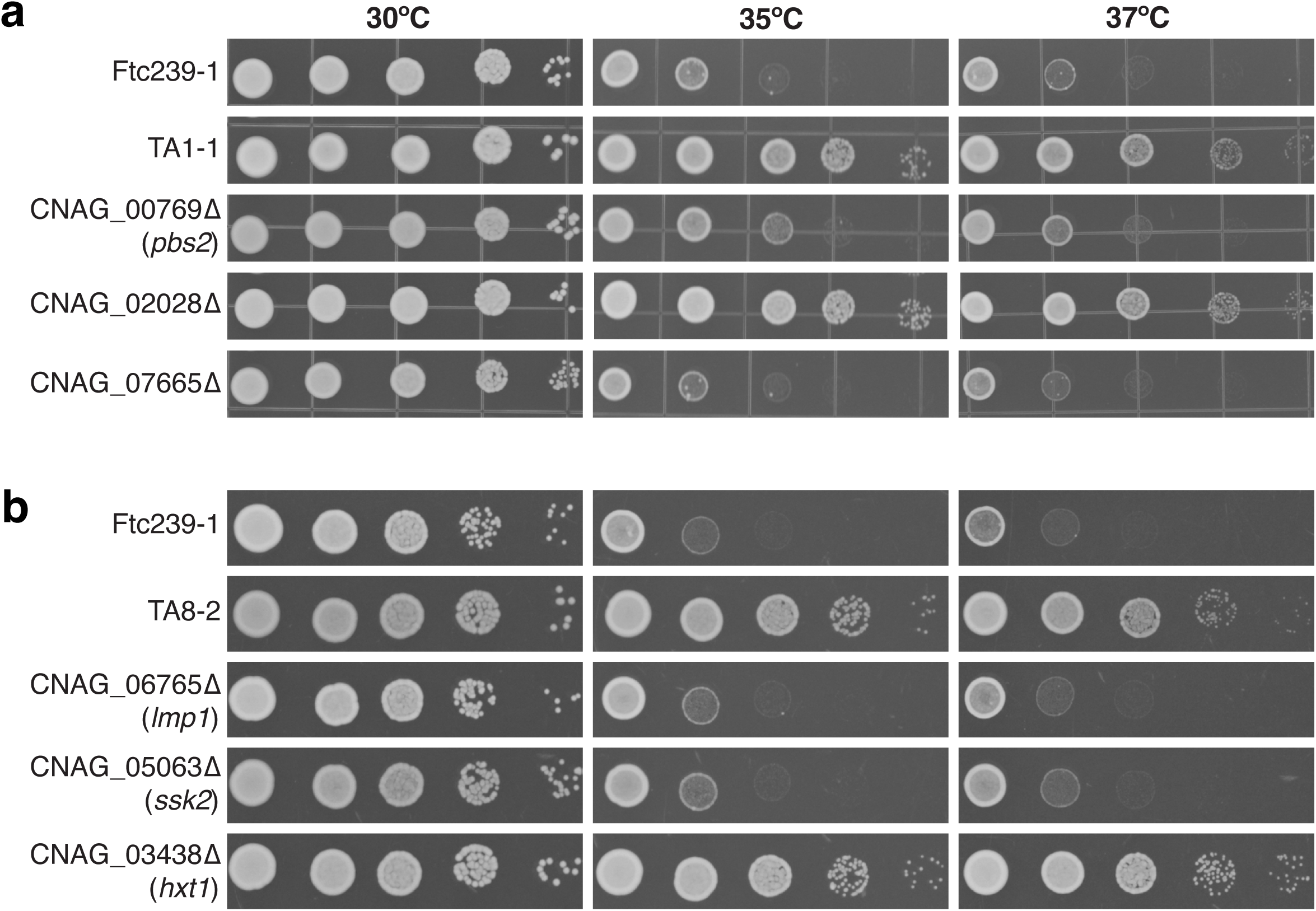
Deletion of the genes *HXT1* or *CNAG_02028* are sufficient for growth at 37°C in the Ftc239-1 background. (a) Comparison of growth across temperatures for the TA1-1 evolved isolate and gene deletion mutants for genes that carry genetic lesions in TA1-1. (b) Comparison of growth of the TA8-2 isolate compared to deletion mutants for the genes that carry coding sequence mutations in TA8-2. All images show serial dilutions plated on YPD and incubated at the indicated temperatures.

## Discussion

Understanding how environmentally derived pathogenic microbes adapt to changes in their environment, such as increasing temperatures, is of critical importance as climate change advances. In this study, we employed experimental evolution to investigate the ability of *Cryptococcus neoformans*, an important human fungal pathogen, to adapt to a high temperature environment and how this alters consequential phenotypes.

We used an environmentally isolated *C. neoformans* strain, Ftc239-1, which exhibited attenuated growth under normal laboratory culture conditions (30°C) and severe growth defects at human body temperature (37°C). During serial passaging at increasing temperatures, however, this strain was able to rapidly adapt, gaining the ability to grow better at high temperature (35°C) than the parental strain after a very brief evolutionary timecourse (Figure 2). The exact timing of when adaptive mutations arose in each population remains to be seen but will further elucidate both the evolutionary trajectory of each independently evolved population and how rapidly this new trait emerges in response to even gradual temperature shifts. Our results also show that the growth advantage of our evolved strains was more pronounced at 37°C, which was outside the range of temperatures used during serial passaging. This suggests that adaptation to modest temperature increases may affect growth at high temperatures more broadly. But how far does this growth advantage extend? Probing whether there is an upper temperature limit to the fitness advantage in these thermally adapted strains is an interesting and important line of future inquiry.

Another critical question raised is whether adaptation to one form of stress (i.e. temperature) confers resistance to other forms of stress and/or alters other infection-relevant traits. Our data show that thermoadapted strains display a number of altered cellular features: from widespread mucoidy and alterations in low-nutrient growth to changes in cell size variation and antifungal drug resistance. Strikingly, our data characterizing antifungal drug resistance across isolates from all populations of our evolution experiment suggest that rising environmental temperatures may not only impact the prevalence of species capable of establishing human infection but may also affect the treatment strategies for such infections (Figure 4a and 4b). Many of our isolates displayed increases in resistance to the common antifungal, fluconazole, as assessed through both MIC determinations and exposure to constant high dose fluconazole. Although it remains to be seen whether this fluconazole resistance is directly linked to thermoadaptation, this result was unexpected and highlights the ease with which fungi can gain resistance to our limited therapeutic options—even in the absence of direct exposure to these drugs. However, increased drug resistance was not a universal feature of our evolved isolates; some, such as TA8-2 displayed increased sensitivity to antifungal drugs, further underscoring the importance of studies examining the interplay between stress resistance, generally, and thermal adaptation.

Through sequencing of two populations, we identified multiple, non-overlapping genetic lesions which likely give rise to the diverse phenotypes we observe in our evolved isolates. Among these, we identified a role for a predicted kinase, *CNAG_02028*, as well as the hexose transporter *HXT1* in promoting thermotolerance in the TA1 and TA8 populations respectively (Figure 5). In both cases, the mutations we recovered in these genes are missense mutations and not clear loss of function alleles; however, the similarities in growth between the full deletion mutants and our evolved clonal isolates hint that these missense mutations are likely to cause loss of protein function. Neither of these genes have any reported role in nor regulation by high-temperature growth in *C. neoformans.* Although sugar metabolism and, in particular, the trehalose pathway in fungi, have been shown to be necessary for surviving high temperature stress, it is particularly intriguing that mutation of just a single hexose transporter—among the dozens encoded in the *C. neoformans* genome—can cause such a dramatic phenotype^26,33^. The exact mechanisms through which *HXT1* participates in thermotolerance and whether there is a link to nutrient conditions remain important outstanding questions.

Beyond the *HXT1* and *CNAG_02028* mutations, we hypothesize that other mutations in our evolved isolates reflect one of several options. They could be passenger mutations that were present when these high fitness impact mutations emerged or they could play more subtle roles in adaptation to the stressful conditions of the experiment including both temperature and other conditions, such as nutrient availability. Future work tracking the emergence of these mutations over the course of the evolution experiment will help shed light on this question. Collectively, our genetic data paired with the diversity of mutations and pathways identified by sampling just two of our eight adapted populations supports the hypothesis that there may be many routes by which this species can evolve thermotolerance.

Our work here begins a foray into the study of how our changing world will alter the pathogens that infect us. As our findings suggest, host body temperature may not represent as severe a barrier to *C. neoformans* and other fungal infections as environmental temperatures increase. This work, therefore, provides a vital foundation to illuminate the consequences a changing world will have for the health of future generations.

## Materials and Methods

### Strains and growth conditions

All strains were stored long term at −80°C in growth media—either yeast extract peptone dextrose (YPD) or yeast nitrogen base (YNB)—supplemented with 20% glycerol. Strains were inoculated by streaking out onto YPD plates and incubated for 2–3 days at 30°C before use in experiments. A full list of strains used can be found in File S1.

### Serial passaging of *C. neoformans* at increasing temperatures

For the first passage, an overnight culture of Ftc239-1 was inoculated from a single colony and grown in YPD with shaking for 16 hours at 30°C. This culture was centrifuged at 3000 × g for 5 minutes and washed twice with autoclaved MilliQ water and resuspended in 5 mL of water before the OD was measured. Cells were normalized to an OD of 0.05 (∼4.04 x 10^6^ cells/ml) in 5 mL of either YPD or YNB with 2% added glucose (4 replicate cultures of each). The remaining starting inoculum culture was mixed 1:1 with 40% glycerol and aliquots were frozen at −80°C.

The cultures were incubated at 30°C with shaking at 225 rpm for 48 hours. After 48 hours, the OD of each individual culture was measured and re-diluted to an OD of 0.05 in 5 mL of the appropriate growth media. Passaging proceeded as above from this point. The temperature was increased once the majority (>75%) of the cultures had reached three to four doublings in the 48 hour culture period. Incubation temperature was increased by 1°C from 30°C to 33°C and 0.5°C from 33°C to 35°C. Aliquots from each individual culture were frozen down as above at the start and end of each temperature, as well as every other passage. Following the final passage at 35°C, multiple aliquots were frozen. Samples from each population were also serially diluted and plated onto YPD plates for CFU determination. For all of the populations except for TA1, serial dilution plates had >50 colonies. Five individual colonies from each of these plates were selected randomly, picked into 5 mL of YPD, grown overnight, and frozen for use in phenotyping experiments (Figure S1, Tables S1 and S2). For population TA1, we only recovered 2 colonies after serial dilution and one was a bacterial contaminant (see discussion below), so the only recovered *C. neoformans* isolate was saved as above. Collectively, this gave us 36 clonal isolates from our eight evolved populations for further phenotyping.

Though the experimental passaging was carried out based on OD_600_ measurements, the actual amount of yeast passaged may not have been adequately reflected by the OD. In the course of our characterization of experimentally evolved strains, we determined that both of the populations discussed in depth here and likely all of the populations were contaminated with *Staphylococcus hominis*, which absorbs at the same OD. This means that passage bottlenecks were more severe than intended. However, this contamination did not affect our ability to isolate thermally adapted fungal clones. Further, our data suggest that the bacterial contaminant has not significantly impacted the thermal adaptation of the *Cryptococcus*.

### Yeast spot assays

*C. neoformans* strains of interest were grown overnight in YPD at 30°C, subcultured to an OD of 0.2 and grown to mid-log phase (OD_600_=0.8-1). The cultures were then centrifuged at 3000 × g for 5 minutes, washed twice with autoclaved MilliQ water, and resuspended in 5 mL of MilliQ water. The OD_600_ of each culture was measured, and the OD was adjusted to OD 1. Ten-fold serial dilutions of each culture were then made down to OD 0.0001. 2.5 microliters of each dilution was spotted onto desired media: YPD, YNB, or YPD plates with the indicated concentrations of fluconazole. Spot plates were incubated at the desired temperature for a total of 72 hours. Plates were imaged on Gel Doc XR+ (BioRad) or using a homemade imaging setup after 48 and 72 hours. Unless otherwise indicated, representative images from spot assays show growth after 48 hours.

### Strain construction and molecular biology

All genetic manipulations were done using CRISPR-mediated gene editing and the TRACE system following previously published protocols^34,35^. Briefly, gRNAs were generated using guide-specific primer and amplification from pBHM2329 plasmid. *CAS9* was amplified for transformation from the pBHM2403 plasmid.

For insertion of the hygromycin B cassette as the Safe Haven 2 locus (SH2), we used a VNB lineage-specific guide targeting this locus. A repair template plasmid carrying the ∼1 kb of sequence immediately 5’ and 3’ to this gRNA sequence from the H99 genome was constructed using HiFi DNA Master Mix (NEB Cat#E2621) with a standard HYG^R^ cassette inserted between the two flanks. Repair templates were then PCR amplified from this plasmid for CRISPR transformations. Gene deletion repair templates were generated following previously published methods. Specifically, the 1000bp of sequence immediately 5’ to the start codon and 3’ to the stop codon were cloned into the pUC19 vector backbone flanking the standard HYG^R^ cassette. These plasmids were sequence verified and then used as template for CRISPR repair template amplification. DNA concentrations in all CRISPR transformations were as follows: 250 ng Cas9, 100 ng gRNA, and 2-2.5 µg repair template.

### Transformant selection and genotyping

Following electroporation of CRISPR reagents, strains were plated onto selective media (YPD+HYG) and allowed to grow for 2-3 days at 30C. Individual colonies were then patched onto non-selective YPD media for 2-3 passages before being patched back onto selective media to identify stable genomic integrants.

Colonies were genotyped by colony PCR following previously published protocols^36^. Genomic sequences were amplified from gDNA extracted from candidate colonies and integration of the drug cassette at the SH2 locus or at each individual gene locus was confirmed by Sanger sequencing. PCRs were also performed to ensure that no strains had integrated the gRNA or *CAS9* constructs. All primer sequences for generating CRISPR reagents and genotyping mutants are listed in File S2.

### Competition assays

For competition experiments, *C. neoformans* strains of interest were grown at 30°C overnight in YPD. For experiments done in YPD, the OD was measured for each test strain and for a parental Ftc239-1 strain carrying a HYG-resistance cassette at the SH2 locus. Cell suspensions at a concentration of ∼4.04 x 10^6^ cells/mL were made for each individual strain. Cell suspensions were mixed 1:1 (test strain:labeled parental) in a total of 40 mL YPD media before adding 5 mL to individual culture tubes. Each competition was performed in biological triplicate for each temperature (30°C or 35°C). CFUs for the initial mixtures were determined by serially diluting the suspensions and plating onto both YPD and YPD+HYG plates. Competition cultures were grown for 24 hours at 225 rpm at their respective temperatures. After 24 hours, serial dilutions were plated onto nonselective and selective media plates as above. Competitive Index here is calculated as: 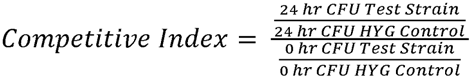

For experiments done in YNB, overnight YPD cultures were washed twice and resuspended in 5 mL of MilliQ water before setting up the experiment as above just in the alternative YNB media.

### Fluconazole MIC determinations

Overnight cultures of *C. neoformans* strains were adjusted to a density of ∼5 x 10^6^ cells/mL and were spread onto either YPD-agar or RPMI-agar medium with L-glutamine and without sodium bicarbonate. Fluconazole Etest strips (bioMérieux) were placed on the agar surface of one of the replicate plates once the cultures had dried. Plates were incubated at 35°C and imaged after 48 and 72 hours. Only strains with robust growth on control YPD plates were imaged and used to determine MIC values.

### Titan cell induction and capsule assays

Titan cell induction was performed following previously published protocols^37^. Briefly, strains were cultured overnight in YPD at 30°C. Cultures were centrifuged at 3000 × g for 5 minutes, washed three times in PBS, and resuspended in 3 mL of PBS. The cells were diluted to ∼5 x 10^3^ cells/ml in 4 mL of serum-free RPMI 1640 media. One milliliter of each strain dilution was plated into three replicate wells of a 24-well plate. The plate was incubated at 37C with 5% CO_2_ without shaking for 7 days. Cells were then fixed with formaldehyde, stained 1:1 with India Ink, and imaged on a Zeiss Axio Observer 7 microscope at 100x.

Capsule was induced following previously published protocols^38^. *C. neoformans* strains of interest were grown overnight (16-18 hours) in YPD at 30°C. Cultures were centrifuged, washed twice with PBS, and resuspended in 5 mL of PBS. OD_600_ was measured for each strain, and the strains were subcultured to an OD of 0.3 in 4 mL of DMEM media containing 10% FBS, non-essential amino acids, and penicillin/streptomycin. Two milliliter aliquots of each dilution were incubated in a 6-well plate at 37°C with 5% CO_2_. After 48 hours of incubation, the cells were fixed and imaged as above.

For both capsule and titan inductions, cell size measurements for each strain were made manually for at least 50 cells under each condition. Capsule thickness here was calculated as: capsule thickness = (total diameter – cell body diameter)/2.

### Genomic DNA extraction and sequencing

Single colonies or pooled culture stocks were inoculated into 50 mL of YPD and grown overnight, or until the culture was fully saturated, with shaking at 30C. Cells were collected by centrifugation and lyophilized overnight. High-molecular-weight genomic DNA was isolated following the CTAB protocol. Following CTAB extraction, samples were further purified using the ProNex Size-Selective Purification System (Promega). Concentrations were measured by Qubit dsDNA broad range assay. For Nanopore sequencing of Ftc239-1, one microgram of HMW gDNA was prepared for sequencing using the SQK_LSK110 Kit and run on an R9 (FLO-MIN106) flow cell. For Illumina sequencing, samples were sheared with a Covaris S2 Focused-ultrasonicator and libraries were prepared using the New England Biolabs NEBNext Ultra II DNA Library Prep kit (cat#E7634L) with an average insert size of 450 bp. Ten strains were barcoded and pooled, and 150×150 bp paired-end reads were collected on a NovaSeq S4 Flow Cell with the v1.5 Reagent Kit. Illumina library preparation and sequencing was performed by the University of Utah’s High Throughput Genomics core facility at the Huntsman Cancer Institute. Raw sequencing datasets are available through the SRA under the BioProject Accession PRJNA1096649.

### Sequencing analysis

A chromosome scale assembly of the parental Ftc239-1 strain was drafted largely as previously published^36^. Briefly, fast5 files from Nanopore sequencing were re-basecalled using guppy-gpu (v6.4.2) and the resulting fastq files were assembled using Canu (v 2.1.1) with an estimated genome size of 20 megabases and default settings^39^. The Canu assembly had 14 chromosome-scale contigs. Shorter contigs mapping to the mitochondrial genome or repetitive genomic regions were manually trimmed. This assembly was polished once with Nanopore reads using Medaka (1.7.2; https://github.com/nanoporetech/medaka) and then polished and additional five times with Illumina reads from the parental strain using pilon (v1.24)^40^. The mitochondrial genome was assembled from Illumina reads using GetOrganelle (v1.7.5.3) with the fungus_mt setting and confirmed by comparison to the trimmed contigs assembled by Canu^41^. This mitochondrial genome was manually added to the assembly. Chromosomes were named based on comparison to other reference assemblies. The complete draft assembly was annotated *de novo* using funannotate (v1.8.15) and publicly available RNA-seq datasets of *C. neoformans* strains grown under a variety of conditions (SRA Accession Numbers: SRR646263, SRR639772, SRR646268, SRR646265)^42^. *De novo* annotations were functionally annotated by BLAST to the reference (H99) proteome available from Uniprot and annotation files were updated using AGAT (v1.0.0)^43^.

For Illumina sequencing data of isolates from the end of the evolution, adapters were trimmed using Trimmomatic (v0.39) and reads were aligned to our *de novo* assembly using bwa-mem2 (v 2.2.1)^44,45^. Duplicates were marked and read groups added with Picard and the resulting files were sorted and indexed with SAMtools^46^. Variants were called using sorted BAM files with FreeBayes (v0.9.21.7) with the following options: -p 1 --min-alternate-fraction 0.8 -- standard-filters^47^. The predicted impacts of called variants were assessed using SnpEff using our *de novo* assembly as reference^48^. Called variants were also cross-checked by analysis with breseq (v0.37.1) with settings altered to reject indels and variants predicted in regions of long homopolymer runs (options: --consensus-reject-indel-homopolymer-length 3 --consensus-reject-surrounding-homopolymer-length 3)^49^. All sequencing analysis was performed using resources through the University of Utah Center for High Performance Computing.

## Data Availability Statement

Strains generated by this study are available upon request. File S1 contains a list of all strains used in this work. Sequencing data are available under the NCBI BioProject Accession PRJNA1096649. Draft genome assembly and annotations for the Ftc239-1 parental isolate have been uploaded to the GSA Figshare portal.

## Acknowledgements and Funding

We thank John Perfect for providing the environmental isolates used in this study. This work was funded by a Helen Hay Whitney Foundation postdoctoral fellowship and Boston College startup funds to ZAH, a Burroughs Wellcome Fund Investigator in the Pathogenesis of Infectious Disease award to NCE, and an NIH R35 (R35GM1 34936) awarded to NCE.

## Author Contributions

Conceptualization: M.J.W.S., Z.A.H.

Methodology: M.J.W.S., L.A.F., M.R., Z.A.H.

Software: Z.A.H.

Validation: M.J.W.S., Z.A.H.

Formal Analysis: M.J.W.S., L.A.F., M.R., J.T.L., Z.A.H.

Investigation: M.J.W.S., L.A.F., M.R., J.T.L., Z.A.H.

Data Curation: M.J.W.S., L.A.F., M.R., J.T.L., Z.A.H.

Writing-Original Draft: M.J.W.S., Z.A.H.

Writing-Review and Editing: M.J.W.S., Z.A.H., N.C.E.

Visualization: M.J.W.S., Z.A.H.

Supervision: Z.A.H., N.C.E.

Funding acquisition: N.C.E., Z.A.H.

## Supplemental Figures and Tables

**Figure S1.**
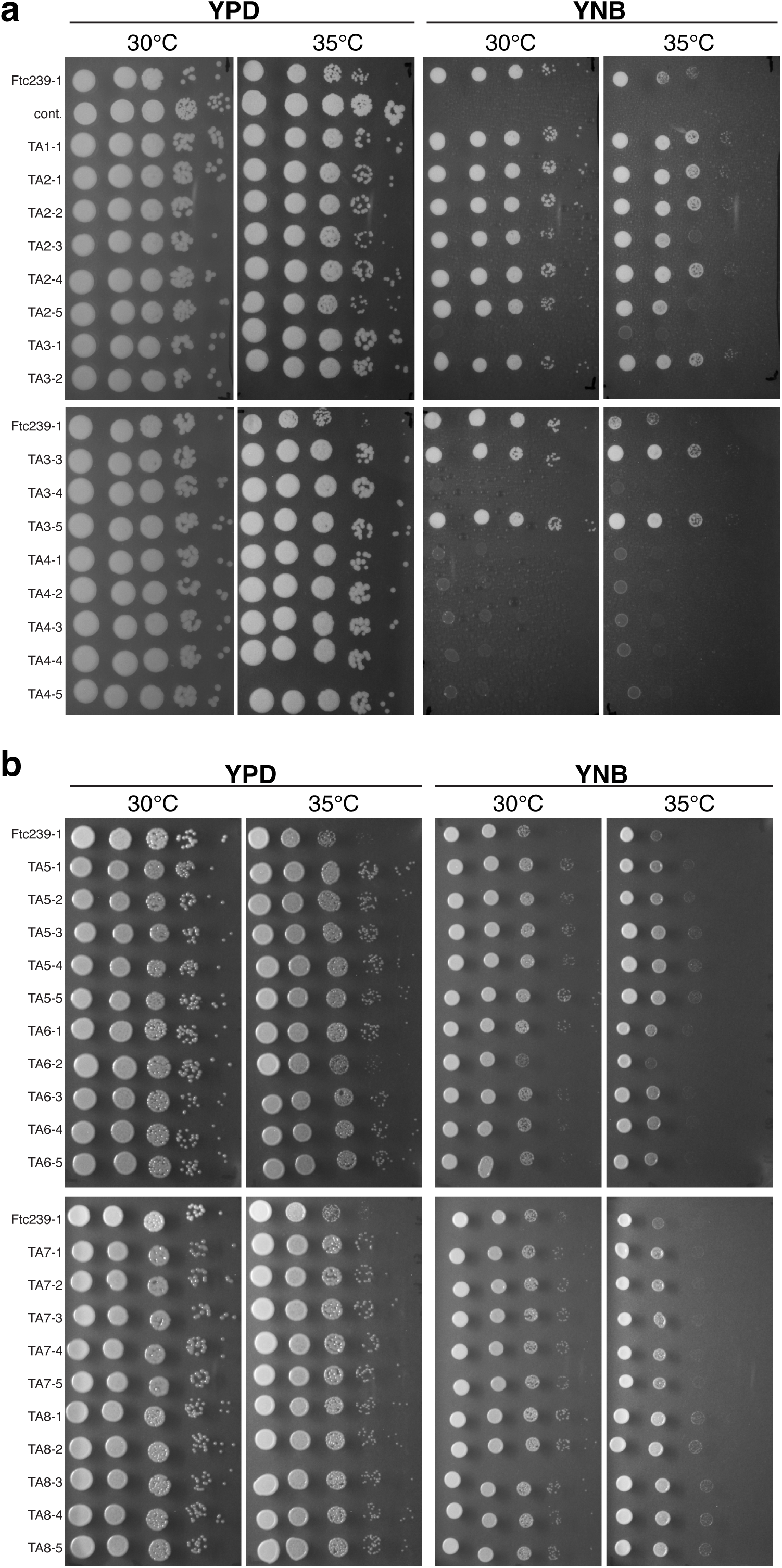
Phenotypic variation of *C. neoformans* parental and evolved isolates. (a and b) Growth of *C. neoformans* parental strain and YPD (A) and YNB (B) evolved isolates on YPD (left) or YNB (right) agar plates at 30 and 35°C.

**Figure S2.**
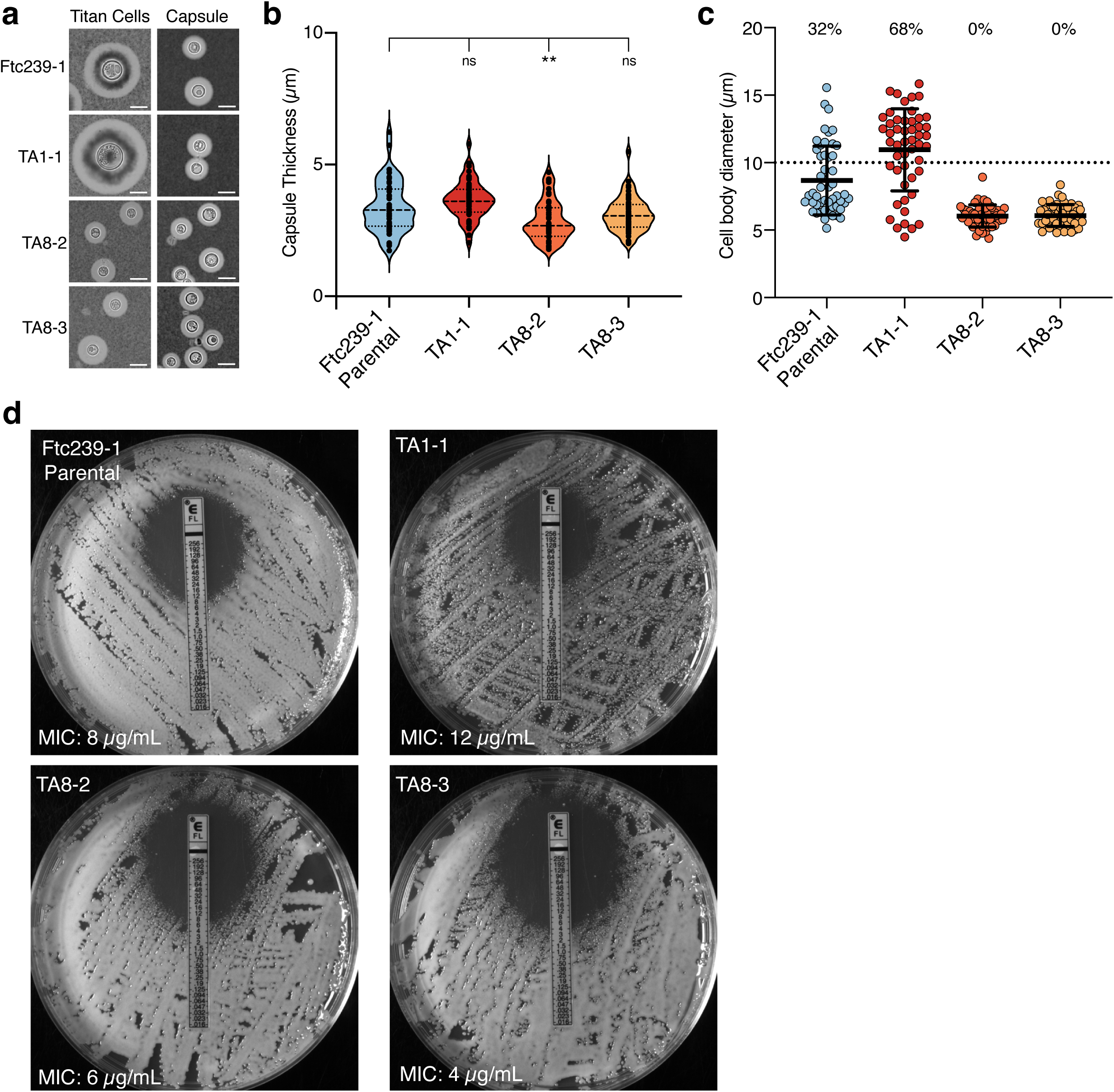
Evolved isolates from two populations display changes in titan cell formation and antifungal resistance. (a) Representative images from cells in titan-inducing (left) and capsule-inducing (right) conditions for parental and evolved strains. Images from different conditions are at the same scale with all scale bars representing 10 µm. (b) Measurements of capsule thickness from cells grown under capsule inducing conditions. Capsule thicknesses were calculated as indicated in Materials and Methods. Median and quartiles are indicated with dashed lines for cells from one representative experiment (c) Measurements of cell body diameter from cells grown under titan inducing conditions. Percentages above each strain indicate the percent of cells with a diameter >10 µm (dotted line). Error bars show mean ± SD for one representative experiment. (d) Fluconazole minimum inhibitory concentrations (MICs) for the parental and evolved isolates on RPMI agar plates with E-test strips. For (b) and (c), 50 cells were measured per strain. Significance was assessed in comparison to the parental Ftc239-1 strain by ordinary one-way ANOVA with Dunnett’s multiple comparisons test.

**Figure S3.**
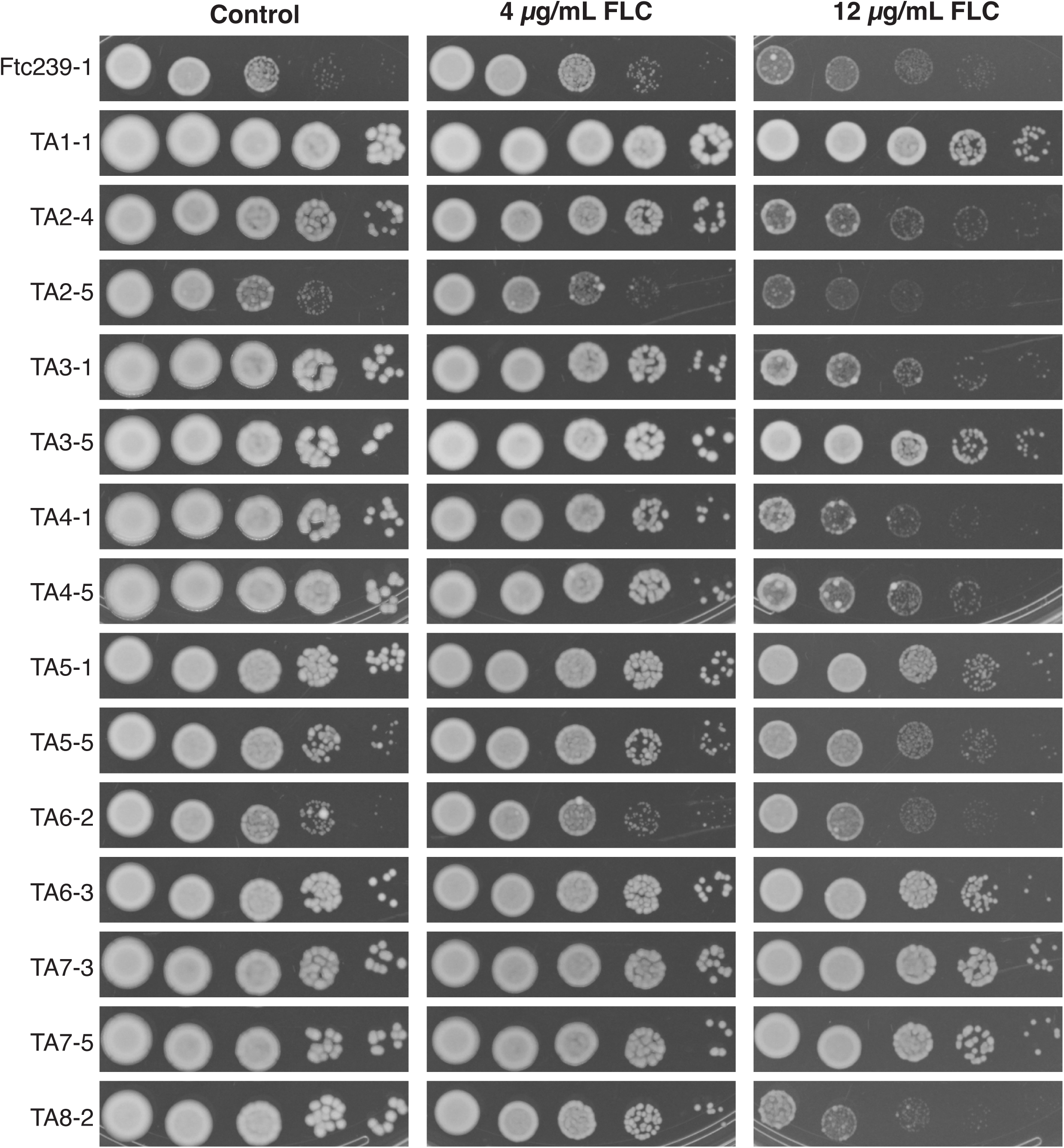
Evolved isolates from many independent populations show enhanced resistance to fluconazole. Growth phenotypes of 14 experimentally evolved isolates on control YPD plates (left) or plates containing the indicated concentrations of fluconazole (right). All plates were incubated at 35°C and images show 72h of growth. Images for strains shown in Figure 4c are identical in this panel and included to allow for comparisons across all of the tested isolates.

**Table S1.**
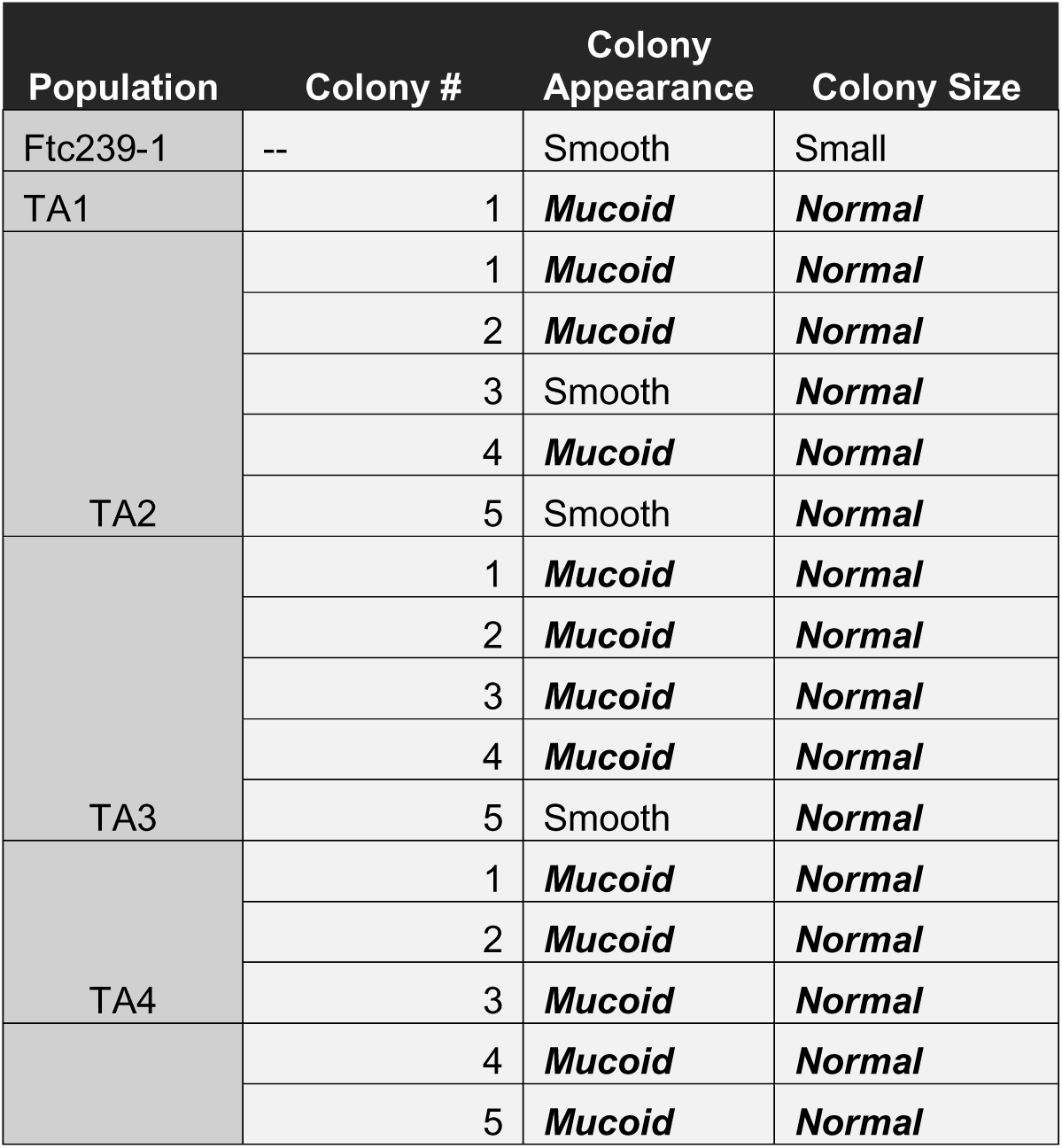
Colony phenotypes of individual colony isolates of the YPD-passaged populations (TA1-TA4). Isolates discussed in this study are indicated by darkened background.

**Table S2.**
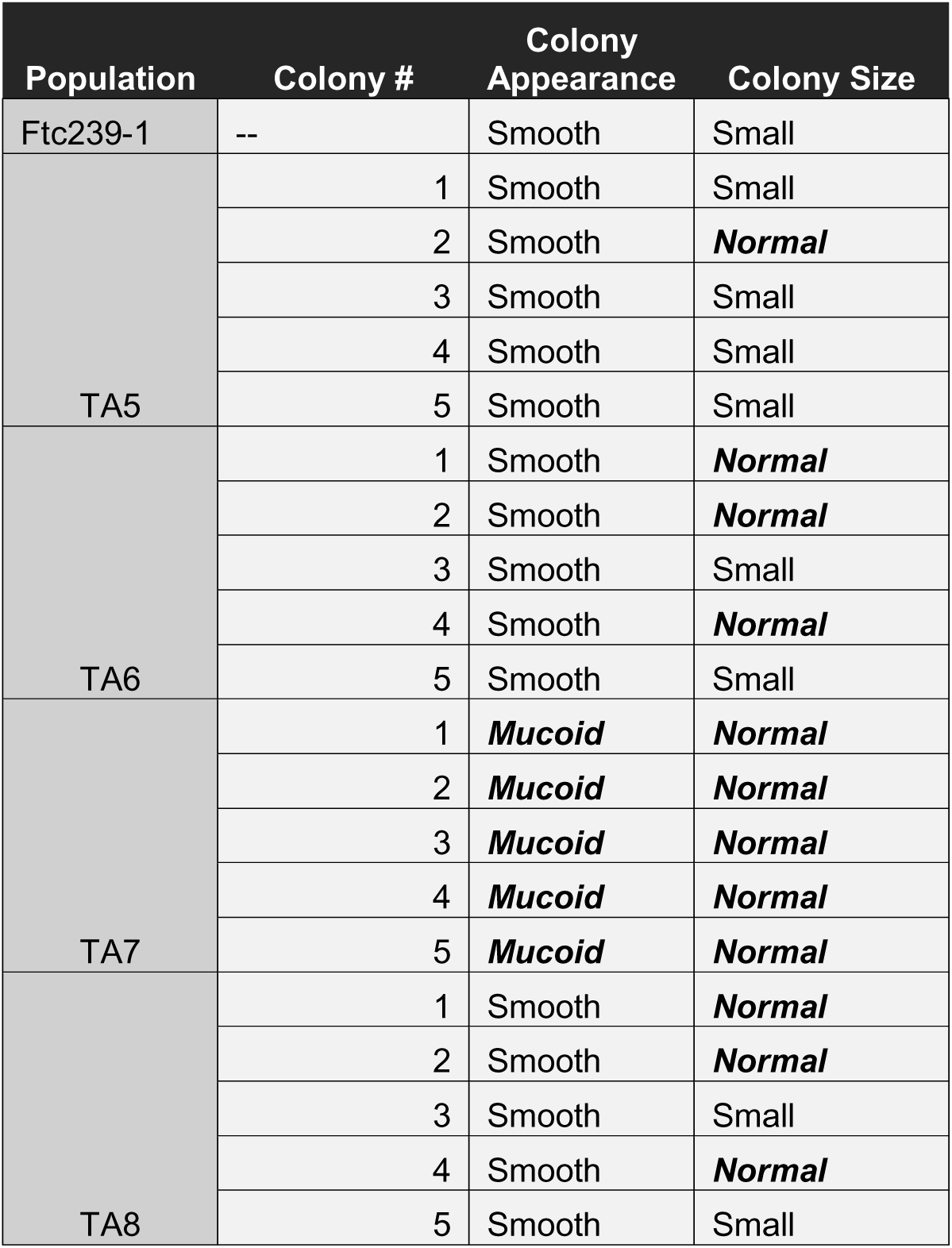
Colony phenotypes of individual colony isolates of the YNB-passaged populations (TA5-TA8). Isolates discussed in this study are indicated by darkened backgrounds.

## Supplemental Files

**File S1.** List of strains used in this study.

**File S2.** List of primers used in this study.

